## Supplementary material for "Quantum theory of a potential biological magnetic field sensor: radical pair mechanism in flavin adenine dinucleotide biradicals"

Supplementary Fig. 1 (following Fig. 3 in the paper) This 2D plot shows $\Phi^{S(S)}$ for different magnetic fields ($B_0$) and distance (r). Singlet fraction yield: $\Phi^{S(S)} = k_s \int_0^\infty Tr\left[\hat{P}^S \hat{\rho}(t)\right] dt$ (initial state: singlet state)

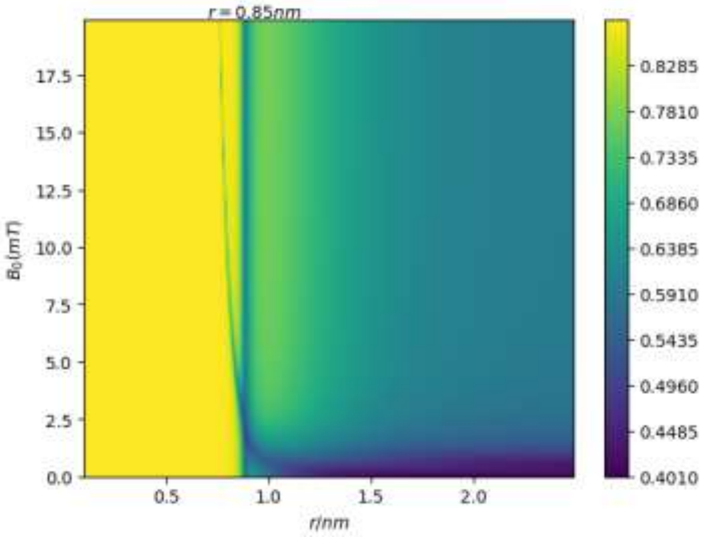


Fig. 1: Singlet yield of a radical pair system with initial singlet state.

Supplementary Fig. 2 (following Fig. 5 in the paper) The original results on the distance between centers of mass of the Adenine and isoalloxazine ring in FAD, calculated using GROMACS. The average of these graphs is illustrated in main text.

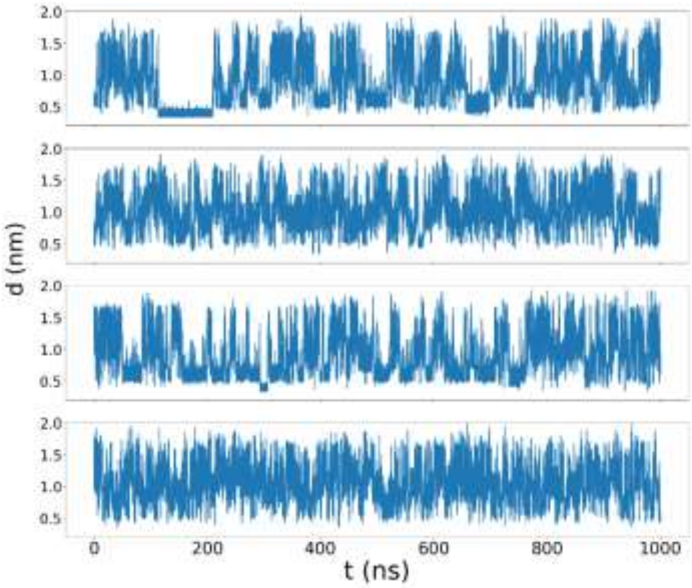


Fig. 2: Distance between centers of mass of the Adenine and isoalloxazine ring. Results obtained from MD simulation with GROMACS.

Supplementary Fig. 3 (following Fig. 7 in the paper) As mentioned in paper, the off-diagonal terms of density operator ($\rho(t)$) are negligible.

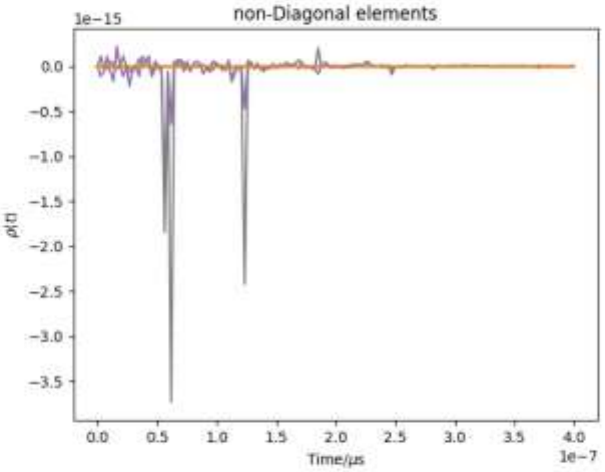


(a) r = 0.5 nm

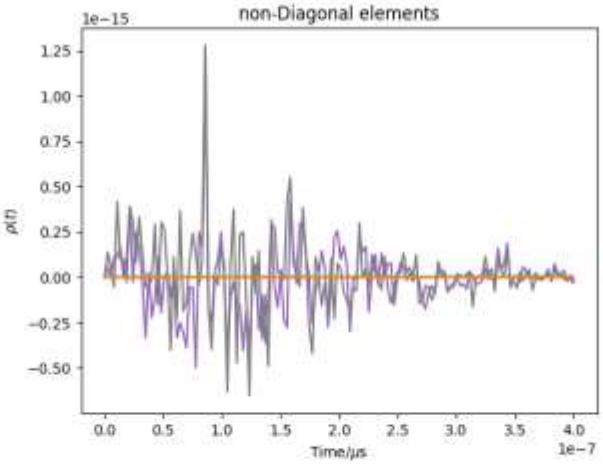


(b) r = 1.5 nm

Fig. 3: Time evolution of the off-diagonal term of density operator for two different distances between radicals.

Supplementary Fig. 4 (following Fig. 10 in the paper) As discussed in paper, ignoring dipole-dipole interaction does not change the overall conclusion about sensitivity of FAD photochemistry to magnetic field.

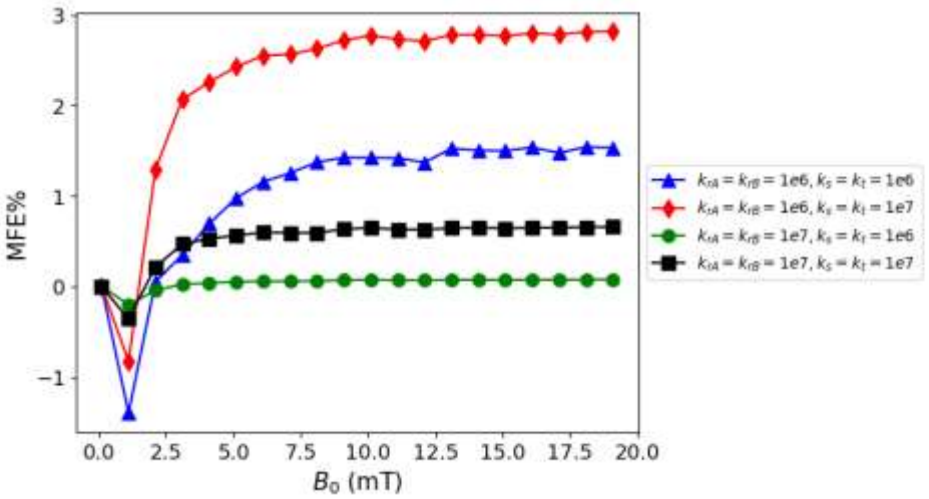


Fig. 4: MFEs of FAD when dipole-dipole interaction is ignored, for different relaxation and reaction rates.

Supplementary Fig. 5. These curves are showing the effect of different HFCC on the MFE of FAD, including the strongest HFCC for flavin ($|a| = 0.8029$) [1].

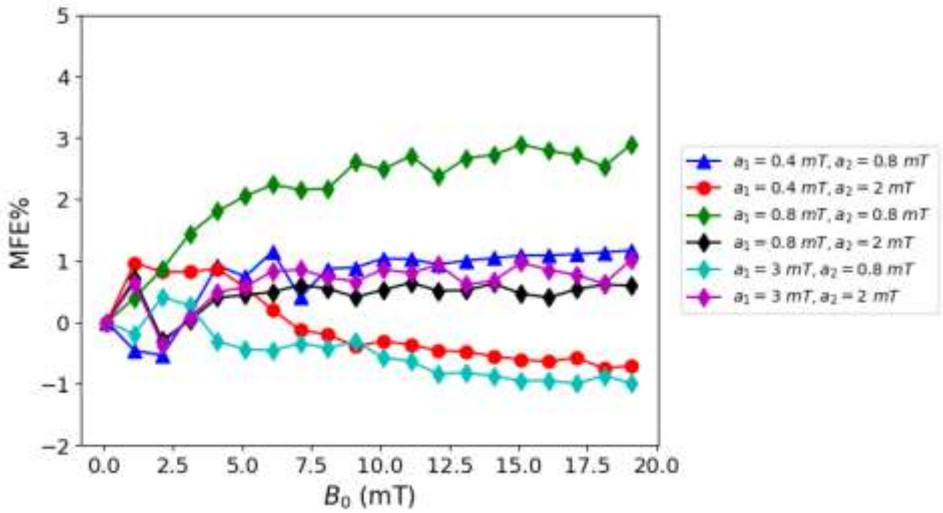


Fig. 5: MFEs of FAD for different HFCC of each radical. The relaxation and recombination rates are $10^6\ s^{-1}$.
